# Evaluation of N^6^-adenine DNA-immunoprecipitation-based genomic profiling in eukaryotes

**DOI:** 10.1101/2022.03.02.482749

**Authors:** Brian M. Debo, Ben Mallory, Andrew B. Stergachis

## Abstract

The detection of low-abundance DNA N^6^-methyladenine (DNA-m6A) remains challenging, limiting our understanding of this novel base in eukaryotes. To address this, we introduce an approach for systematically validating the selectivity and sensitivity of antibody-based DNA-m6A methods, revealing most commercial antibodies as poorly selective towards DNA-m6A. Finally, using a validated highly selective anti-DNA-m6A antibody we expose distinct patterns of DNA-m6A in *C. reinhardtii*, *A. thaliana*, and *D. melanogaster*.

## INTRODUCTION

Endogenous m6A-DNA is a widespread DNA modification in bacterial genomes(Ratel et al. 2006a) and some unicellular eukaryotes (Fu et al. 2015; Hattman et al. 1978), and more recently has been described in multicellular eukaryotes such as *Arabidopsis thaliana* (Liang et al. 2018), *Caenorhabditis elegans* (Greer et al. 2015), *Drosophila melanogaster* (Zhang et al. 2015), *Mus musculus* (Wu et al. 2016; Koziol et al. 2016; Li et al. 2020), *Homo sapiens* (Wu et al. 2016; Xie et al. 2018), and others (Luo and He 2017). The level of DNA-m6A in eukaryotes has been reported to vary from as low as 6 ppm (i.e., 6 m6A-modified bases per million adenines) in humans and mice (Wu et al. 2016) to as high as 8,000 ppm in *Tetrahymena pyriformis*(Hattman et al. 1978), and can vary by over 10-fold between different tissues within the same organism (Zhang et al. 2015; Li et al. 2020). However, the biological role of DNA-m6A in multicellular eukaryotes remains largely undefined as the accuracy of genome-wide DNA-m6A methodologies for the detection of low-abundance DNA-m6A is disputed (Lentini et al. 2018; Douvlataniotis et al. 2020; Schiffers et al. 2017; Ratel et al. 2006b; O’Brown et al. 2019; Kong et al. 2022). Specifically, although antibodies targeting modified bases have been a workhorse for the study of other base modifications, results from anti-DNA-m6A antibodies have thus far proven to be unreliable (O’Brown et al. 2019), with microbial contaminants thought to be a major contributor to these false measurements (Kong et al. 2022; Douvlataniotis et al. 2020).

## RESULTS

To directly evaluate N^6^-adenine DNA-immunoprecipitation-based detection methods, we sought to develop a straightforward approach for systematically testing the selectivity and sensitivity of anti-DNA-m6A antibodies for the global and locus-specific detection of DNA-m6A. To accomplish this, we leveraged a recently described approach for generating complex genomic DNA (gDNA) libraries with titratable amounts of locus-specific DNA-m6A (Stergachis et al. 2020). Specifically, we used gDNA from eukaryotic cells lacking significant endogenous m6A-DNA (negative control), as well as gDNA isolated after treating nuclei from these cells with a non-specific m6A-methyltransferase (m6A-MTase) (positive control). We then tested ten commercial anti-DNA-m6A antibodies against these samples to systematically quantify the sensitivity and selectivity of these anti-m6A antibodies (**Fig. 1a**). We demonstrated that commonly used cell culture lines, such as drosophila S2-DRSC, and human K562 cells can be used for generating the positive and negative control samples, as these cells lack appreciable endogenous DNA-m6A (**Fig. 1b** and **Supplementary Fig. 2**), and treating nuclei from these cells with an m6A-MTase results in the methylation of ~14% of all adenines across various sequence contexts (Stergachis et al. 2020) (i.e., DNA-m6A amount of 140,000 ppm) (**Supplementary Fig. 1**).

**Figure 1.**
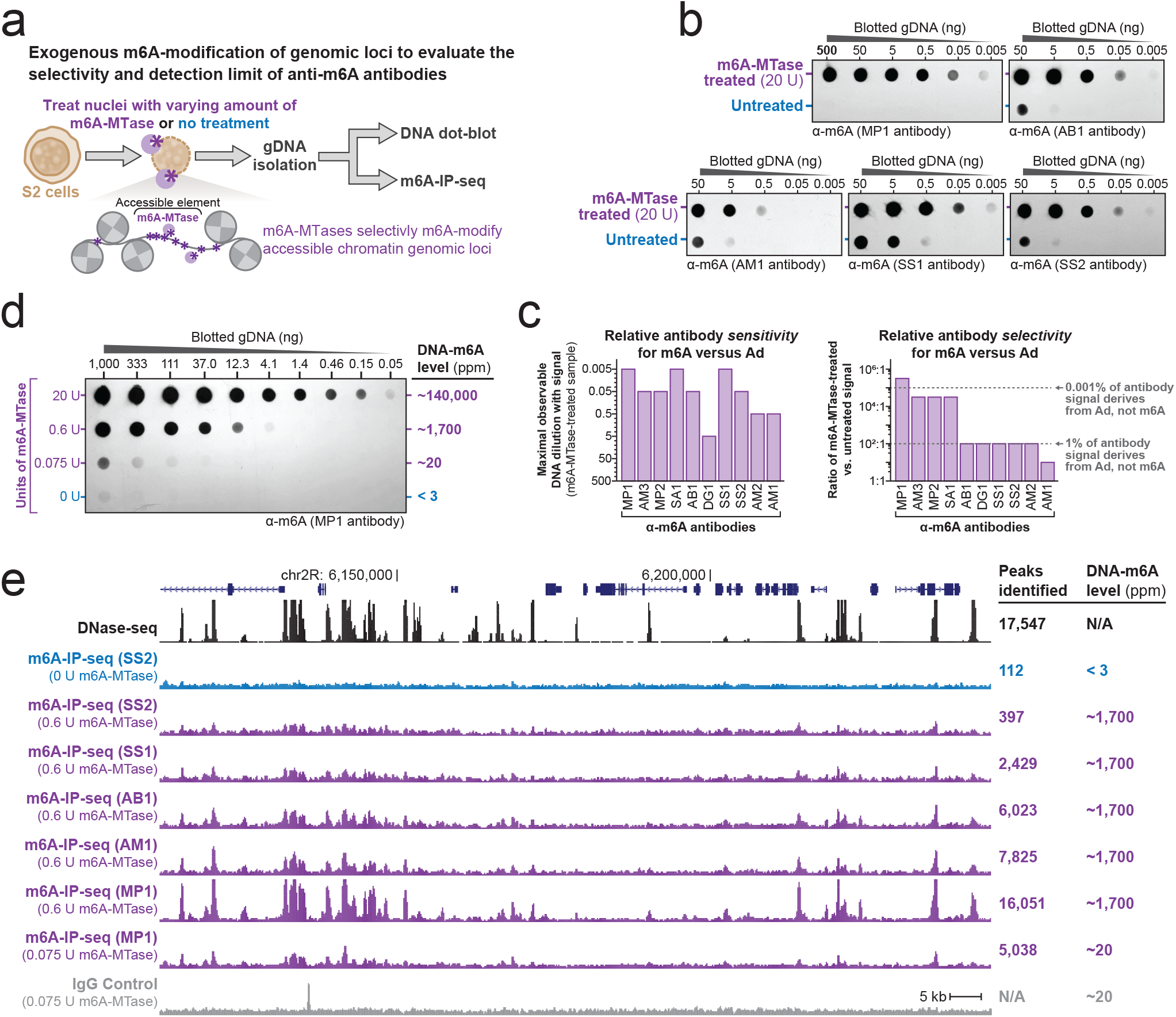
Sensitivity and selectivity of anti-DNA-m6A antibodies. (a) Schematic for assaying anti-DNA-m6A antibody selectivity and sensitivity using gDNA isolated from cell nuclei exogenously methylated with a non-specific DNA m6A-MTase. (b) DNA dot blots using five separate anti-DNA-m6A antibodies against gDNA from untreated S2 cells versus gDNA from S2 cells treated with m6A-MTase. (c) Bar chart quantifying the relative sensitivity (left) and selectivity (right) of each anti-DNA-m6A antibody towards DNA-m6A, as measured using DNA dot-blots. (d) DNA dot blot and table demonstrating the relative amount of m6A-DNA after treatment of S2 cell nuclei with increasing amounts of a non-specific DNA-m6A-MTase. (e) Genomic locus comparing the relationship between DNaseI-seq and m6A-IP-seq signal. m6A-IP-seq performed using five separate antibodies on untreated S2 cell gDNA, or gDNA from S2 cell nuclei treated with 0.6 U or 0.075 U of m6A-MTase. Signal from IgG antibody control also displayed. Y-axis is identical for all m6A-IP-seq and DNase I-seq experiments.

We found that all ten antibodies detected DNA-m6A, albeit with sensitivities that differed by over two orders of magnitude (**Figs. 1b-c** and **Supplementary Fig. 2**). However, the selectivity of these antibodies for DNA-m6A, as opposed to unmethylated Ad, varied quite dramatically between the ten antibodies. For example, whereas the MP1 antibody demonstrated over a 100,000-fold selectivity towards DNA-m6A, the AM1 antibody was only 10-fold selective towards DNA-m6A as opposed to unmethylated Ad (**Figs. 1b-c** and **Supplementary Fig. 2**).

To determine the limit of detection for the MP1 antibody, we titrated the amount of m6A-MTase used to generate the positive control sample – showing that the MP1 antibody can detect DNA-m6A levels as low as ~3 ppm (**Fig. 1d**). In contrast, most of the other anti-DNA-m6A antibodies could only quantify DNA-m6A levels down to ~1,400 ppm before the DNA-m6A signal became indistinguishable from off target antibody recognition of unmethylated adenines – precluding the use of these antibodies for quantifying endogenous DNA-m6A levels in essentially all multicellular eukaryotic organisms (Luo and He 2017).

Next, we sought to evaluate the sensitivity of antibody-based DNA-m6A methods for identifying endogenous DNA-m6A-modified genomic loci using these exogenously methylated positive control samples. Specifically, we applied DNA-m6A immunoprecipitation followed by massively parallel sequencing (m6A-IP-seq) to gDNA isolated from S2 cell nuclei treated with 0.6 units of m6A-MTase (**Fig. 1a**), as this treatment results in a global DNA-m6A level of ~1,700 ppm (**Fig. 1d**) and the selective deposition of DNA-m6A at ~17,000 accessible chromatin elements genome-wide in a pattern mirroring that of DNaseI-seq (Stergachis et al. 2020). For this approach, we only tested five of the antibodies with varying degrees of sensitivity and specificity for DNA-m6A. Notably, only the MP1 antibody accurately detected nearly all exogenously DNA-m6A-methylated sites genome-wide, with the other four antibodies detecting only between 2-45% of the exogenously methylated sites (**Fig. 1e** and **Supplementary Fig. 3**), suggesting that these four antibodies exhibit both poor selectivity and sensitivity for the detection of m6A-modified bases and genomic loci.

To test the detection limit of DNA-m6A-antibodies for identifying modified genomic loci, we applied m6A-IP-seq to gDNA isolated from S2 cell nuclei treated with only 0.075 units of m6A-MTase. This treatment results in a global DNA-m6A level of ~20 ppm (**Fig. 1d**), which is similar to endogenous DNA-m6A levels in multicellular eukaryotes (Luo and He 2017). Although m6A-IP-seq signal using the MP1 antibody was markedly reduced in this sample, we still identified ~29% of the genomic loci exogenously modified with DNA-m6A (**Fig. 1e**), demonstrating that appropriately sensitive and selective anti-DNA-m6A antibodies can detect low-abundance DNA-m6A modified loci.

Next, using a highly selective and sensitive anti-DNA-m6A antibody (e.g., MP1), we reexamined endogenous DNA-m6A levels across three diverse eukaryotic organisms reported to contain endogenous DNA-m6A at similar levels – *C. reinhardtii* (Fu et al. 2015; Hattman et al. 1978), *A. thaliana* cauline leaves (Liang et al. 2018), and *D. melanogaster* 45-minute embryos (Zhang et al. 2015) (**Fig. 2a**). Overall, we found that whereas *C. reinhardtii* demonstrated endogenous DNA-m6A levels consistent with prior reports, both the *A. thaliana* and *D. melanogaster* samples demonstrated endogenous DNA-m6A levels 30-50-fold lower than previously reported (Liang et al. 2018; Zhang et al. 2015; Luo and He 2017) (**Figs. 2a-b**), establishing a more limited presence of endogenous DNA-m6A in *A. thaliana* cauline leaves and *D. melanogaster* 45-minute embryos.

**Figure 2.**
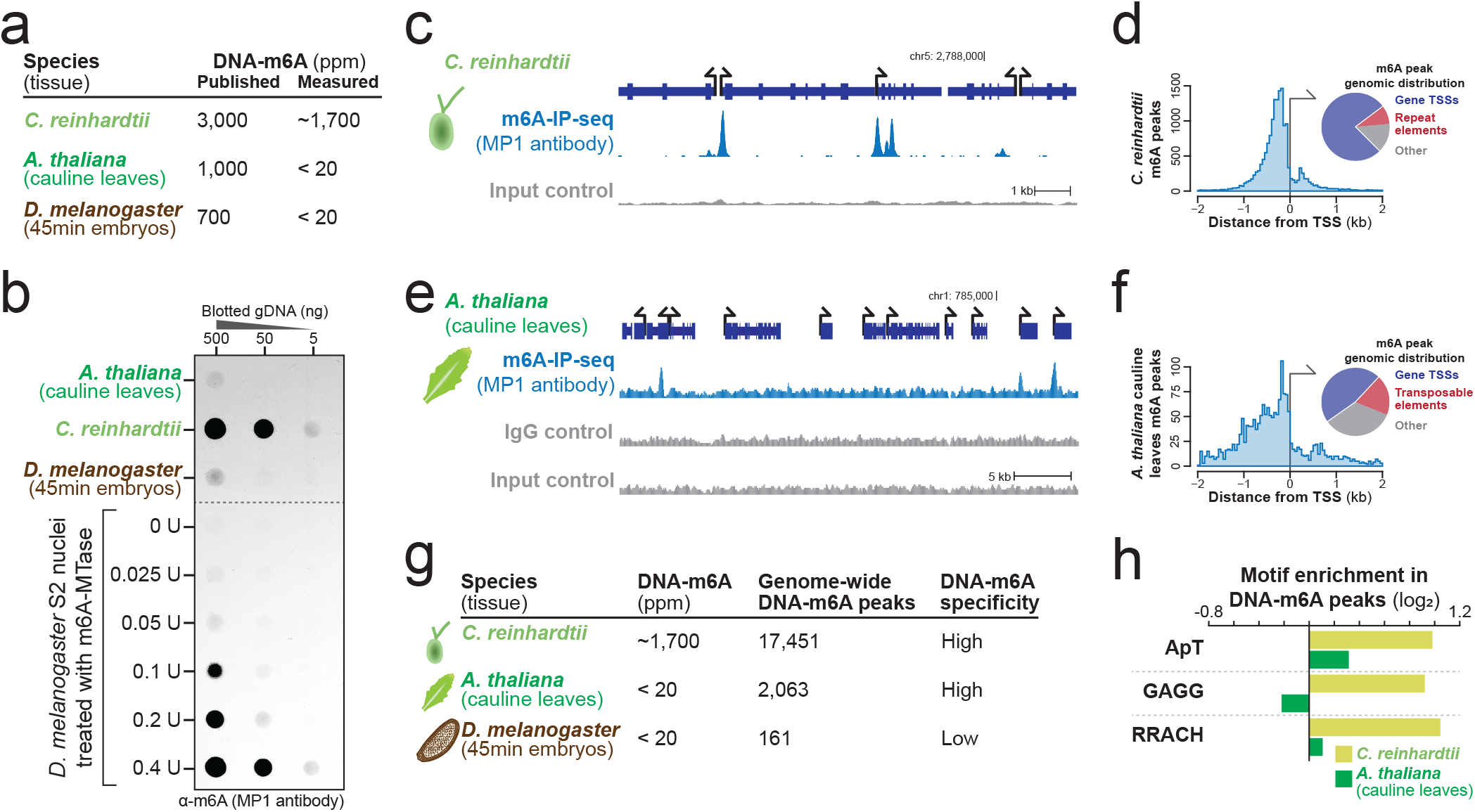
Detection of endogenous DNA-m6A across three non-mammalian eukaryotes. (a) Table showing the published and observed amount of DNA-m6A in samples from three eukaryotes. (b) DNA dot blot quantifying the relative amount of m6A-DNA in samples from three eukaryotes, as well as from S2 cell nuclei treated with increasing amounts of m6A-MTase. (c) Genomic locus from *C. reinhardtii* comparing the relationship between m6A-IP-seq signal and input control signal. Y-axis is identical for both experiments. (d) (left) Histogram and (right) pie chart demonstrating the distribution of identified *C. reinhardtii* DNA-m6A peaks relative to TSSs and repetitive elements. (e) Genomic locus from *A. thaliana* comparing the relationship between m6A-IP-seq signal and input and IgG control signal. Y-axis is identical for all experiments. (f) (left) Histogram and (right) pie chart demonstrating the distribution of identified *A. thaliana* DNA-m6A peaks relative to TSSs and repetitive elements. (g) Table showing the observed amount of DNA-m6A in samples from three eukaryotes, as well as the number of genomic loci identified with DNA-m6A signal. *Indicates possible false positive calls. (h) Motif enrichment relative to shuffled sequences of three previously reported eukaryotic m6A-MTase motifs in DNA-m6A peaks from *C. reinhardtii* and *A. thaliana*.

Performing m6A-IP-seq on *C. reinhardtii* cells with the MP1 antibody revealed 17,451 DNA-m6A sites genome wide, mirroring previously published data (**Figs. 2c-d** and **Supplementary Fig. 4**) (Fu et al. 2015). Notably, m6A-IP-seq performed using the SS1 antibody, which is the antibody used in the prior *C. reinhardtii* study (Fu et al. 2015), resulted in a high background signal (**Supplementary Fig. 4**), suggesting that some anti-DNA-m6A antibodies may exhibit large lot-to-lot variability, thus explaining some of the poor reproducibility associated with anti-DNA-m6A antibodies (Xie et al. 2018; Wu et al. 2016).

Performing m6A-IP-seq on *A. thaliana* cauline leaves with the MP1 antibody identified 2,063 endogenous DNA-m6A sites genome-wide. These peaks were enriched at gene and transposable element transcriptional start sites (TSSs) (**Figs. 2e-f**), consistent with prior reports (Liang et al. 2018). In contrast, performing m6A-IP-seq on *D. melanogaster* 45-minute embryos identified only 161 DNA-m6A peaks genome-wide, many of which overlapped m6A-IP-seq peaks in S2 cells (**Fig. 2g** and **Supplementary Fig. 5**). These findings indicate that the minimal amount of DNA-m6A we detected in *D. melanogaster* embryos is either not sitespecific within the mappable portion of the genome, or results from residual contamination (**Fig. 2a-b**).

Finally, using our high-quality maps of endogenous DNA-m6A loci in *A. thaliana* and *C. reinhardtii,* we sought to determine whether a similar DNA-m6A MTase is likely mediating DNA-m6A-modified loci in these organisms. Three distinct classes of DNA-m6A MTase motifs have been described to date: the ApT motif, which is methylated by the MTA1c complex in ciliates (Beh et al. 2019); the GAGG motif, which is methylated by the DAMT-1 enzyme in *C. elegans* (Greer et al. 2015); and the RRACH motif, which is methylated in ssDNA by the METTL3-METTL14 complex in *H. sapiens* (Woodcock et al. 2019). Notably, all three of these motifs appeared to be enriched in the *C. reinhardtii* DNA-m6A-modified loci (**Fig. 2h**), indicating that multiple m6A-MTases may contribute to the establishment of DNA-m6A-modified loci in *C. reinhardtii.* In contrast, none of these motifs were appreciably enriched in *A. thaliana* DNA-m6A-modified loci. Rather, *A. thaliana* DNA-m6A-modified loci demonstrated a separate highly enriched motif (**Supplementary Fig. 6**), indicating that distinct pathways are likely generating DNA-m6A in *A. thaliana* and *C. reinhardtii*.

## DISCUSSION

In conclusion, we demonstrate a robust approach for systematically evaluating the selectivity and sensitivity of anti-DNA-m6A antibodies using a controlled system that enables the titratable DNA-m6A methylation of thousands of loci genome-wide across various sequence contexts. Using this approach, we demonstrate that most commercially available anti-DNA-m6A antibodies exhibit poor selectivity and sensitivity towards DNA-m6A, which includes one of the most commonly used anti-DNA-m6A antibodies(Xie et al. 2018; Hao et al. 2020; Fu et al. 2015; Liang et al. 2018; Greer et al. 2015; Wu et al. 2016; Koziol et al. 2016). These findings highlight the role of poor antibody selectivity, in addition to bacterial and RNA contamination(Kong et al. 2022), to prior reports of abundant DNA-m6A across diverse eukaryotic organisms, and potentially hold broader implications beyond DNA-m6A genomic profiling as many commercial anti-m6A antibodies are used interchangeably with DNA and RNA-m6A methodologies. Furthermore, using a highly selective anti-DNA-m6A antibody we expose distinct biological features of endogenous DNA-m6A in *C. reinhardtii* and *A. thaliana,* and demonstrate a lack of site-specific DNA-m6A in *D. melanogaster* 45 min embryos. Given the low level of DNA-m6A in eukaryotes, our approach provides a tractable system for accurately detecting DNA-m6A modified loci and for continually reassessing the sensitivity and selectivity of anti-DNA-m6A antibodies.

## METHODS

### Production of non-specific DNA m6A-MTase

The non-specific DNA-m6A-MTase Hia5 was produced as previously described (Stergachis et al. 2020). Of note, the method presented in this manuscript can also be performed using other non-specific DNA-m6A-MTases, including the commercially available EcoGII (NEB catalog # M0603S). We have previously demonstrated that at least five non-specific DNA-m6A-MTase can be used interchangeably (Stergachis et al. 2020).

### Treatment of S2 cell nuclei with m6A-MTAse and gDNA isolation

Drosophila S2-DRSC cells were grown and isolated as previously described (Stergachis et al. 2020). Specifically, three million S2 cells were pelleted at 250 x g for 5 minutes then resuspended in 60 μl of Buffer A (15 mM Tris, pH 8.0; 15 mM NaCl; 60 mM KCl; 1mM EDTA, pH 8.0; 0.5 mM EGTA, pH 8.0; 0.5 mM Spermidine). 60 μl of cold 2X Lysis buffer (0.1 % IGEPAL CA-630 in Buffer A) was added to each sample and mixed by gentle flicking then kept on ice for 10 minutes. Samples were pelleted at 4°C for 5 min at 350 x g and the supernatant was removed. Nuclei pellets were gently resuspended with wide bore pipette tips in 57.5 μl Buffer A and moved to a 37°C thermocycler. 1.5 μl SAM (0.8 mM final) and Hia5 m6A-MTase diluted in Buffer A were added, then carefully mixed by pipetting with wide bore tips. The specific reaction conditions are as follows: 20 U Hia5 for 10min; 0.6 U Hia5 for 10min; 0.4 U Hia5 for 10min; 0.2 U Hia5 for 10min; 0.1 U Hia5 for 10min; 0.075 U Hia5 for 10min; 0.05 U Hia5 for 10min; and 0.025 U Hia5 for 10min. The reactions were incubated at 37°C then stopped with 3 μl of 20% SDS (1% final) and transferred to new 1.5 mL microfuge tubes. The sample volumes were increased by adding 130 μl Buffer A and an additional 7 μl of 20% SDS. All samples were mixed with 3 μl Proteinase K (NEB P8107S) and incubated for 1 hour at 50°C. The DNA was purified by adding 200 μl (1:1) of phenol:chloroform:isoamyl alcohol 25:24:1 (saturated with 10mM Tris, pH 8.0, 1mM EDTA) then mixed by vigorous tube inversions and incubated for 10 min at room temperature (RT). Extractions were centrifuged for 10 min at 17,900 x g and the upper aqueous phase was transferred to new microfuge tubes and the DNA precipitated by adding 0.5 μl GlycoBlue Coprecipitant (Invitrogen AM9515), 0.1 volumes of 3M sodium acetate, pH 5.2, and 2.5 volumes ice cold 100% ethanol. The DNA was pelleted by centrifuging at 20,000 g for 10 min at 4°C and washed by repeating the centrifugation with 1 mL of ice cold 70% ethanol. The tubes were inverted over a tube rack and air dried for 15 minutes before resuspension in 200 μl of Buffer A. All samples were mixed with 3 μl of RNase A (Invitrogen AM2271) and incubated for 1 hour at 37°C. The samples subsequently underwent an additional phenol:chloroform:isoamyl alcohol purification, and residual phenol was removed with a second extraction by adding 200 μl chloroform:isoamyl alcohol 24:1 to all samples and repeating the extraction procedure. The DNA ethanol precipitation, washing, pelleting, and drying were repeated before resuspension in 54 μl of 10 mM Tris, pH 7.5.

### Treatment of K562 cell nuclei with m6A-MTAse and gDNA isolation

Human K562 cells were grown and isolated as previously described(Stergachis et al. 2020). Reaction conditions were similar to those described for S2 cells above, with the following differences: (a) 1 million K562 cells were used per reaction; (2) 2X Lysis buffer consisted of 0.05 % IGEPAL CA-630 in Buffer A; (c) 400 U Hia5 was used; and (d) Hia5 reaction was performed for 10min at 25°C prior to being stopped with the addition of SDS.

### *A. thaliana* cauline leaf genomic DNA extraction

Cauline leaves were collected from three separate flowering col-0 Arabidopsis thaliana plants and combined together. Two biological replicates were obtained from plants grown 3 months apart from each other. Approximately 1 g of Cauline leaves was combined with 1.2 mL of Buffer A supplemented with 1% sodium dodecyl sulfate (Sigma L3771), 0.5% polyvinylpyrrolidone (Sigma PVP40) and fresh 1% 2-Mercaptoethanol (Sigma M3148) in a Dounce homogenizer. Tissue was ground using a “B” pestle until homogenous. The lysate was transferred in 200 μl aliquots to 1.5 mL microfuge tubes and all samples were mixed with 3 μl Proteinase K and incubated for 1 hour at 50°C. The DNA was purified by adding 200 μl (1:1) of phenol:chloroform:isoamyl alcohol 25:24:1 (saturated with 10mM Tris, pH 8.0, 1mM EDTA) then mixed by vigorous tube inversions and incubated for 10 min at RT. Extractions were centrifuged for 10 min at 17,900 x g and the upper aqueous phase was transferred to new microfuge tubes and the DNA precipitated by adding 0.5 μl GlycoBlue Coprecipitant (Invitrogen AM9515), 0.1 volumes of 3M sodium acetate, pH 5.2, and 2.5 volumes ice cold 100% ethanol. The DNA was pelleted by centrifuging at 20,000 g for 10 min at 4°C and washed by repeating the centrifugation with 1 mL of ice cold 70% ethanol. The tubes were inverted over a tube rack and air dried for 15 minutes before resuspension in 200 μl of Buffer A. All samples were mixed with 5 μl of RNase A (Invitrogen AM2271) and incubated for 1.5 hours at 37°C. The DNA purification with phenol:chloroform:isoamyl alcohol was repeated and residual phenol was removed with a second extraction by adding 200 μl chloroform:isoamyl alcohol 24:1 to all samples and repeating the extraction procedure. The DNA ethanol precipitation, washing, pelleting, and drying were repeated before resuspension in 25 μl of 10 mM Tris, pH 7.5 and pooling of all samples.

### C. reinhardtii genomic DNA extraction

Chlamydomonas reinhardtii (cc-5155, Chlamydomonas Resource Center) was grown in TAP media until log phase (2-6 × 10^6^ cells/mL) under continuous illumination at 100 μM photons/m^2^/sec. Cells were pelleted, and DNA was extracted from the cell pellet as described above for *A. thaliana* cauline leaves.

### D. melanogaster 45 minute embryo genomic DNA extraction

Drosophila melanogaster early embryos were collected from young (<7 days) male and female wild type flies. Two biological replicates were collected 2 months apart. Approximately 100-200 flies were transferred to large embryo collection cages and fed baker’s yeast paste for three days prior to collection. Fresh 100mm molasses agar plates were exchanged daily. Timed egg lays were used to stage embryos by adding a fresh molasses agar plate with yeast paste to the collection cage for 30 minutes; this was then replaced with a fresh plate for the next collection cycle. The plate was submerged in 1X PBS and the eggs were dislodged with a paint brush then inverted over a funnel into an egg basket and rinsed with additional PBS. The egg basket was then inverted over a 50 mL conical tube and the eggs were collected with PBS and pelleted for 3 min at 300 x g. The PBS was removed and the eggs were transferred with a wide bore pipette tip in approximately 1 mL of PBS to a 1.5 mL microfuge tube and pelleted for 2 min at 500 x g. The PBS was removed and the egg pellet was flash frozen in liquid nitrogen and stored at −80°C. On average, the embryo pellets were frozen 15 minutes after removal of the agar plate from the collection cage yielding embryos between 15 and 45 minutes of development (referred hereafter as 45min embryos). The pellets were transferred to 1.2 mL of Buffer A supplemented with 1% sodium dodecyl sulfate, 0.5% polyvinylpyrrolidone and fresh 1% 2-Mercaptoethanol, and DNA was extracted as described above for *A. thaliana* cauline leaves.

### Antibodies

Primary antibodies used in this paper:

- Rabbit polyclonal anti-N6-methyladenosine antibody (Millipore Sigma ABE572 – RRID:AB_2892213), referred hereafter as “MP1”
- Rabbit polyclonal anti-N6-methyladenosine antibody (Millipore Sigma ABE572-I – RRID:AB_2892214), referred hereafter as “MP2”.
- Mouse monoclonal anti-N6-methyladenosine antibody (Active Motif 61756 – RRID:AB_2793759), referred hereafter as “AM1”
- AbFlex^®^ N6-Methyladenosine (m6A) antibody (rAb) (Active Motif 91261 – RRID:AB_2892216), referred hereafter as “AM2”.
- N6-Methyladenosine (m6A) antibody (pAb) (Active Motif 61495 - RRID:AB_2793658), referred hereafter as “AM3”.
- Rabbit polyclonal anti-N6-methyladenosine antibody (Synaptic System Ab. 202 003 – RRID:AB_2279214), referred hereafter as “SS1”
- Mouse monoclonal anti-N6-methyladenosine antibody (Synaptic System Ab. 202 111 - RRID:AB_2619891), referred hereafter as “SS2”.
- Rabbit polyclonal anti-N6-methyladenosine antibody (Abcam ab151230 – RRID:AB_2753144), referred hereafter as “AB1”
- Anti-N6-Methyladenosine (m6A) antibody, Rabbit monoclonal (Sigma-Aldrich SAB5600251 – RRID:AB_2892215), referred hereafter as “SA1”.
- N6-methyladenosine (m6A) Antibody (Diagenode C15200082-50 – RRID:AB_2892212), referred hereafter as “DG1”.

Secondary antibodies used in this paper:

- Anti-rabbit IgG HRP-linked (Cell Signaling Technology 7074 - RRID:AB_2099233)
- Anti-mouse IgG HRP-linked (Cell Signaling Technology 7076 - RRID:AB_330924)

### DNA-m6A dot blot

DNA sample dilutions were made in a 20X SSC buffer in a 96-well plate followed by denaturation at 95°C for 10 min. Nitrocellulose membrane was wetted in 20X SSC buffer, secured in a HYBRI-DOT Manifold (Life technologies), then washed under vacuum with 20X SSC buffer, followed by the addition of denatured DNA samples to each well. Membrane was then placed on a dry Whatman filter paper and crosslinked with 125 mJoule in a GS Gene Linker UV Chamber (Bio-Rad) using the C-L setting. The membrane was then washed with 20 mLs of 1X TBS-T (10 mM Tris, pH 7.5; 0.25 mM EDTA; 150 mM NaCl; 0.1% TWEEN-20) and blocked in 15 mLs of 1X TBS-T + 5% non-fat dry milk for 1 hour at RT. All primary antibodies were diluted 1:1000 in 10 mLs 1X TBS-T + 5% non-fat dry milk, and primary incubation was performed overnight at 4°C on a slow shaker. The blot was washed 3 times in 20 mLs of 1X TBS-T for 15 minutes total. Secondary antibodies were diluted 1:2000 in 15 mLs 1X TBS-T + 5% non-fat dry milk and incubated with the blot for 1 hour at room temperature. Three washes were performed and the blot was developed with Pierce ECL Plus Western Blotting Substrate (Thermo Scientific 32132) and imaged using film.

### DNA-m6A Immunoprecipitation sequencing (m6A-IP-seq)

Genomic DNA was sheared using a Covaris M220 Focused-Ultrasonicator (peak power: 75.0; duty factor: 15.0%; cycles/burst: 200; duration: 720 seconds; water bath temperature: 20°C) to achieve a fragment length of approximately 100 base pairs. Libraries were constructed from these sheared DNA samples using the NEBNext Ultra II DNA Library Prep Kit for Illumina (NEB E7645S) and NEBNext Multiplex Oligos for Illumina Index Primers Sets 1 and 2 (NEB E7335S & E7500S). In order to increase input DNA above single library limits (1 μg) for samples with the lowest DNA-m6A abundance (D. melanogaster embryos and A. thaliana cauline leaves), duplicate end prep and adaptor ligation reactions were prepared and combined into single immunoprecipitations (including IgG controls). The samples were end repaired by combining 1 μg of DNA in 50 μl of 10 mM Tris, pH 8.0 with 7 μl of End Prep Reaction Buffer and 3 μl of End Prep Enzyme Mix in PCR strip tubes. Samples were mixed and placed on a thermocycler using the end repair program (30 minutes at 20°C, 30 minutes at 65°C, then hold at 4°C). Adaptors were ligated to samples by adding 2.5 μl of the Adaptor for Illumina, followed by 30 μl of the Ligation Master Mix, then 1 μl of the Ligation Enhancer. Samples were incubated for 15 minutes at 20 °C, then 3 μl of USER enzyme was added and incubated for 15 minutes at 37 °C. For cleanup, 116 μl (1.2X) of Agencourt AMPure XP beads (Beckman Coulter A63880) was added to each sample and mixed, followed by 10 minutes of RT incubation and 10 minutes of magnetic separation. The supernatant was discarded and the beads were washed twice on the magnet with 200 μl of fresh 80% ethanol then dried for four minutes. Samples were removed from the magnet and 50 μl of 10 mM Tris, pH 7.5 was added followed by 10 minutes of RT incubation and 10 minutes of magnetic separation. The samples were transferred to new 1.5 mL LoBind tubes, combined with 20 μl of 100 μM blocking oligo (AGATCGGAAGAGCGTC; duplicate reactions are pooled and combined with 40 μl of blocking oligo), and incubated at 95°C for 10 minutes to denature the DNA. Tubes were then transferred to an ice/water slush mix for 10 minutes for rapid cooling. To each sample, 325 μl 0.1X TE buffer (255 μl for pooled samples), and 100 μl 5X IP Buffer (50 mM Tris, pH 7.5; 750 mM NaCl; 0.5% IGEPAL CA-630) were added in addition to 5 μg of anti-N6-methyladenosine antibody (see above for catalog numbers) or Normal Rabbit IgG (Sigma Millipore, NI01-100UG). Samples were incubated for 12 hrs on a rotator at 4°C. 25 μl of Protein A Dynabeads (Invitrogen 10001D) per sample were prepared by removing supernatant after magnetic separation and washing 4 times in 1X IP buffer (10 mM Tris, pH 7.5; 150 mM NaCl; 0.1% IGEPAL CA-630) followed by resuspension in 25 μl 1X IP buffer. The washed Protein A beads were added to each sample and rotated for 4 hours at 4°C. Samples were then washed with 750 μl 1X IP buffer six times, with each wash consisting of 5 minutes of rotation at 4°C. DNA was then eluted by adding 48 μl of Proteinase K Digestion Buffer (20 mM HEPES, pH 7.5; 1 mM EDTA; 0.5% SDS) and 2 μl proteinase K to the beads and incubating at 50°C for 1 hr with 1200 rpm of shaking. Samples were separated on a magnet and the supernatant was transferred to new strip tubes. For cleanup, 90 μl (1.8X) of AMpure XP beads was added to each sample following the same procedure above but with elution in 17 μl 10 mM Tris, pH 7.5 followed by quantification with the Qubit ssDNA Assay Kit (Invitrogen, Q10212). For PCR enrichment, 15 μl of each sample was transferred to a new strip tube and combined with 5 μl of Universal PCR primer, 5 μl of Index primer (both provided in NEBNext Multiplex Oligos for Illumina Index Primers Sets 1 and 2), and 25 μl NEBNext Ultra II Q5 Master Mix (PCR cycling conditions: 98°C for 30 seconds; five to seven cycles of 98°C for 10 seconds then 65°C for 75 seconds; final 65°C incubation for 5 min). For cleanup, 60 μl (1.2X) of AMpure XP beads was added to each sample following the same procedure above, but with elution in 33 μl of 10 mM Tris, pH 7.5. Libraries were quantified using a Qubit dsDNA HS Assay Kit (Q32851) and size distribution was checked on an Agilent Bioanalyzer High Sensitivity DNA chip. Libraries were sequenced using an Illumina HiSeq 4000 to a read depth of ~20 million reads using paired end 76bp read lengths.

### Input control samples

Genomic DNA from S2 cells, A. thaliana cauline leaves, D. melanogaster 45 minute embryos and C. reinhardtii cells were sheared as above and underwent library construction as above. These samples were then amplified by adding the NEBNext Ultra II Q5 Master Mix as above (Program: 98°C of 30 seconds; three cycles of 98°C for 10 seconds then 65°C for 75 seconds; final 65°C incubation for 5 min), and cleaned using 1.2X AMpure XP beads as above. Libraries were sequenced using an Illumina HiSeq 4000 to a read depth of ~20 million reads using paired end 76bp read lengths.

### DNaseI-seq

Previously published DNaseI-seq data was used (Stergachis et al. 2020) (GEO accession number GSE146942).

### Absolute quantification of DNA-m6A levels

The absolute quantity of DNA-m6A in S2 cell nuclei treated with 20 U of Hia5 for 10 min was determined using Pacific Biosciences (PacBio) Circular consensus sequencing. This PacBio data was previously published by our group (Stergachis et al. 2020) (GEO accession number GSE146942). Specifically, we restricted our analysis to reads with 50 or more circular consensus passes, which enables the highly accurate assessment of absolute m6A levels (O’Brown et al. 2019). The methylation profile on each read was determined as previously described (Stergachis et al. 2020), and the average adenine methylation level was then calculated, which was 14%. The absolute quantity of DNA-m6A in other samples was derived by comparing the relative dot-blot signal intensity to that of the 20 U Hia5 sample. Specifically, the dilution of the 20 U Hia5 sample needed to achieve a similar dot-blot intensity to the sample of interest; this was used to calculate the relative amount of DNA-m6A in the sample of interest.

### Identification of DNA-m6A modified genomic loci

Reads were mapped to their respective genome as previously described (Stergachis et al. 2020). Genome builds used were dm6 (*D.melanogaster*), TAIR9 (*A. thaliana*), and creinhardtii_281_v5.0 (*C. reinhardtii*). Signal tracks were generated using BEDOPS (Neph et al. 2012) and signal was normalized to 1 million reads. For the Drosophila S2 samples, DNA-m6A peaks were identified using the MACS2 software (Zhang et al. 2008) using the IgG control sample as the background. For the *D.melanogaster* 45 minute embryo and *A. thaliana* cauline leaf samples, DNA-m6A peaks were identified using the MACS2 software (Zhang et al. 2008) using both the IgG control sample and the input control sample as the background. For the *C. reinhardtii* samples, DNA-m6A peaks were identified using the MACS2 software (Zhang et al. 2008) using the input control sample as the background. A q-value cut-off of 0.01 was used for each sample. Biological replicates were performed for the *D.melanogaster* 45 minute embryo and *A. thaliana* cauline leaf samples, but only a single replicate was used for analyses.

### DNA-m6A loci genomic positions

A. thaliana gene and transposable element transcriptional start sites were identified using the ‘TAIR9_GFF3_genes_transposons.gff’ annotation file downloaded from the Arabidopsis Information Resource (Huala et al. 2001). C. reinhardtii gene transcriptional start sites were identified using the ‘Crein-hardtii_281_v5.5.gene_exons.gff3’ annotation file, and C. reinhardtii repeat elements were identified using the ‘Creinhardtii_281_v5.5.repeatmasked_assembly_v5.0.bed’ annotation file, both of which were downloaded from the Phytozome JGI(Merchant et al. 2007). The position of the center of each DNA-m6A peak to each of these element classes was determined using BEDOPS(Neph et al. 2012).

### DNA-m6A motif enrichment

The ApT motif was derived from (Beh et al. 2019). The GAGG motif was derived from (Greer et al. 2015). The RRACH motif was derived from (Woodcock et al. 2019). Motif enrichments were calculated by comparing the number of motifs identified in DNA-m6A peaks from a given species versus the number of motifs identified in shuffled sequences of DNA-m6A peaks from that same species. Motif instances were identified using FIMO (Grant et al. 2011) with a p-value cutoff of 0.001 for the GAGG and RRACH motifs and 0.01 for the ApT motif. De novo motif discovery using MEME suite (Bailey and Elkan 1994) was performed on *A. thaliana* cauline leaf DNA-m6A peaks. Motif instances within *A. thaliana* cauline leaf and *C. reinhardtii* DNA-m6A peaks were identified using FIMO and a p-value cutoff of 0.0001.

## DATA ACCESS

All sequencing data presented in this manuscript is available at a GEO accession GSE197747.

All code used in this manuscript is described within the methods section and can be made available upon request.

## ACKNOWLEDGEMENT

We thank Aaron Aker for kindly providing *C. reinhardtii* cells. We thank Jennifer Bush for kindly providing *A. thaliana* plants. We thank Richard Binari for kindly providing *D. melanogaster* flies. We thank Eric Haugen and Richard Sandstrom for computational assistance. We thank L. Stirling Churchman and John A. Stamatoyannopoulos for helpful comments on the manuscript. This research was supported by NIH grants DP5-OD029630 (OD) and GM007748 (NIGMS). A.B.S. holds a Career Award for Medical Scientists from the Burroughs Wellcome Fund.

## AUTHOR CONTRIBUTIONS

A.B.S and B.M.D. designed the experiments. A.B.S, B.M.D, and B.M. performed the experiments. A.B.S performed the computational analyses. A.B.S. and B.M.D wrote the manuscript.

## Figure Legends

**Supplementary Figure 1.**
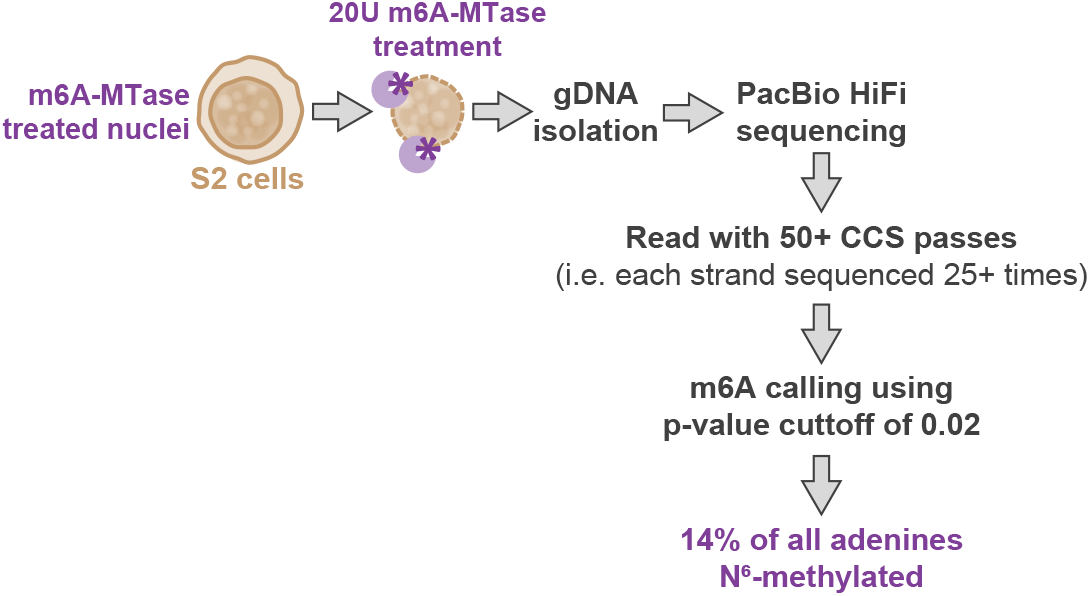
Determining absolute DNA-m6A amount on a highly methyated sample using PacBio single-molecule circular consensus sequencing (CCS). Schematic demonstrating how Drosophila S2 cell nuclei are treated with 20 units of a non-specific m6A-MTase, followed by genomic DNA isolation, DNA shearing to 1-4kb in length and PacBio circular consensus sequencing using a Sequel I platform. DNA-m6A events were identified on a per-molecule basis using a p-value cut off of 0.02, and analysis was restricted to molecules with 50+ circular passes to ensure robust DNA-m6A identification. Overall, this approach identified that 14% of all adenines are marked by DNA-m6A after treatment of S2 cell nuclei with 20 units of a non-specific m6A-MTase.

**Supplementary Figure 2.**
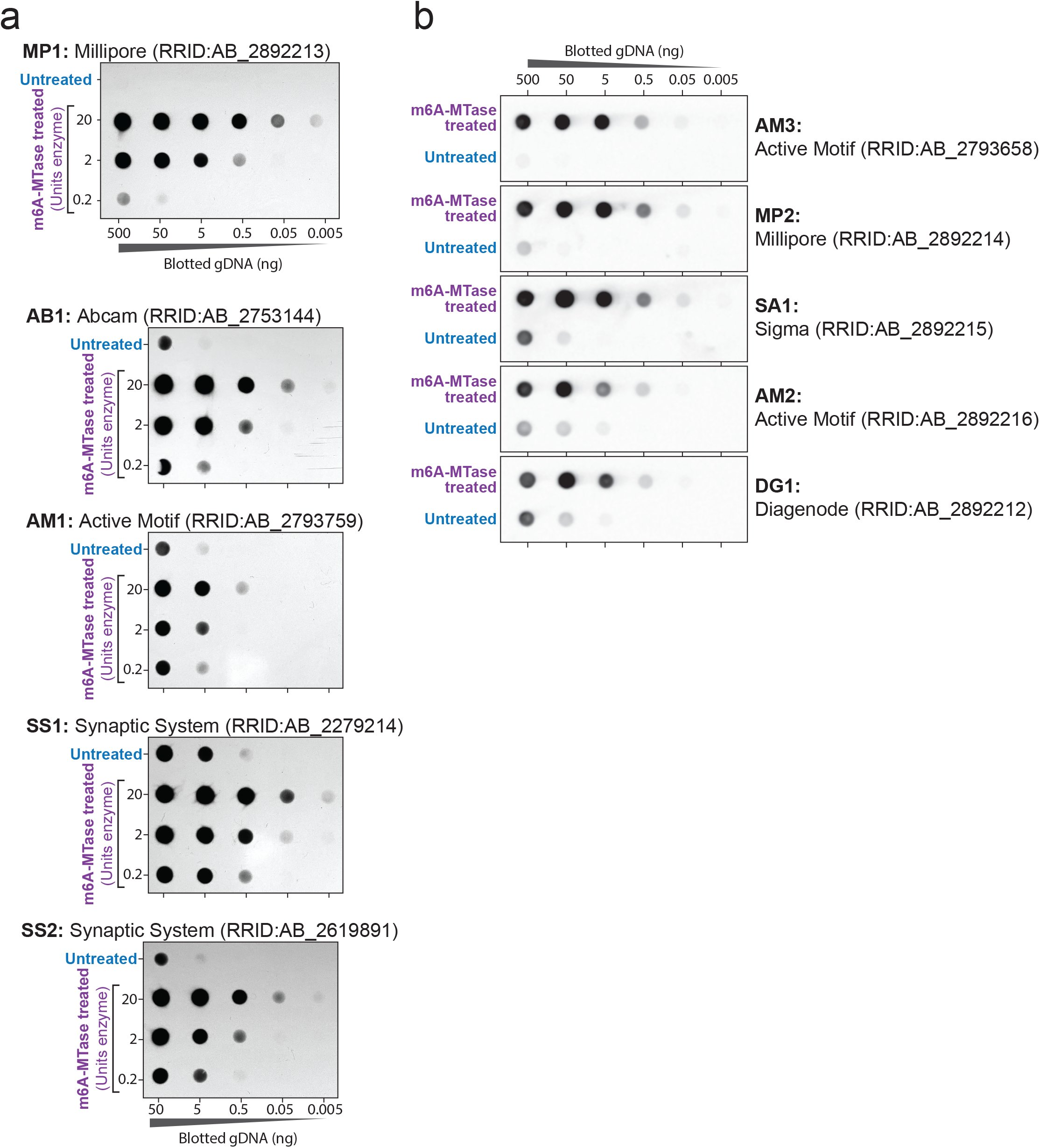
Selectivity of anti-DNA-m6A antibodies towards DNA-m6A. (a) DNA dot blots using 5 separate anti-DNA-m6A antibodies against genomic DNA from untreated S2 cells, versus genomic DNA from S2 nuclei treated with various amounts of a non-specific DNA m6A-MTase. The top two rows of each blot are identical to those shown in Figure 1B. (b) DNA dot blots using 5 separate anti-DNA-m6A antibodies against genomic DNA from untreated K562 cells, versus genomic DNA from K562 nuclei treated with a non-specific DNA m6A-MTase (400U Hia5 for 10min at 25C).

**Supplementary Figure 3.**
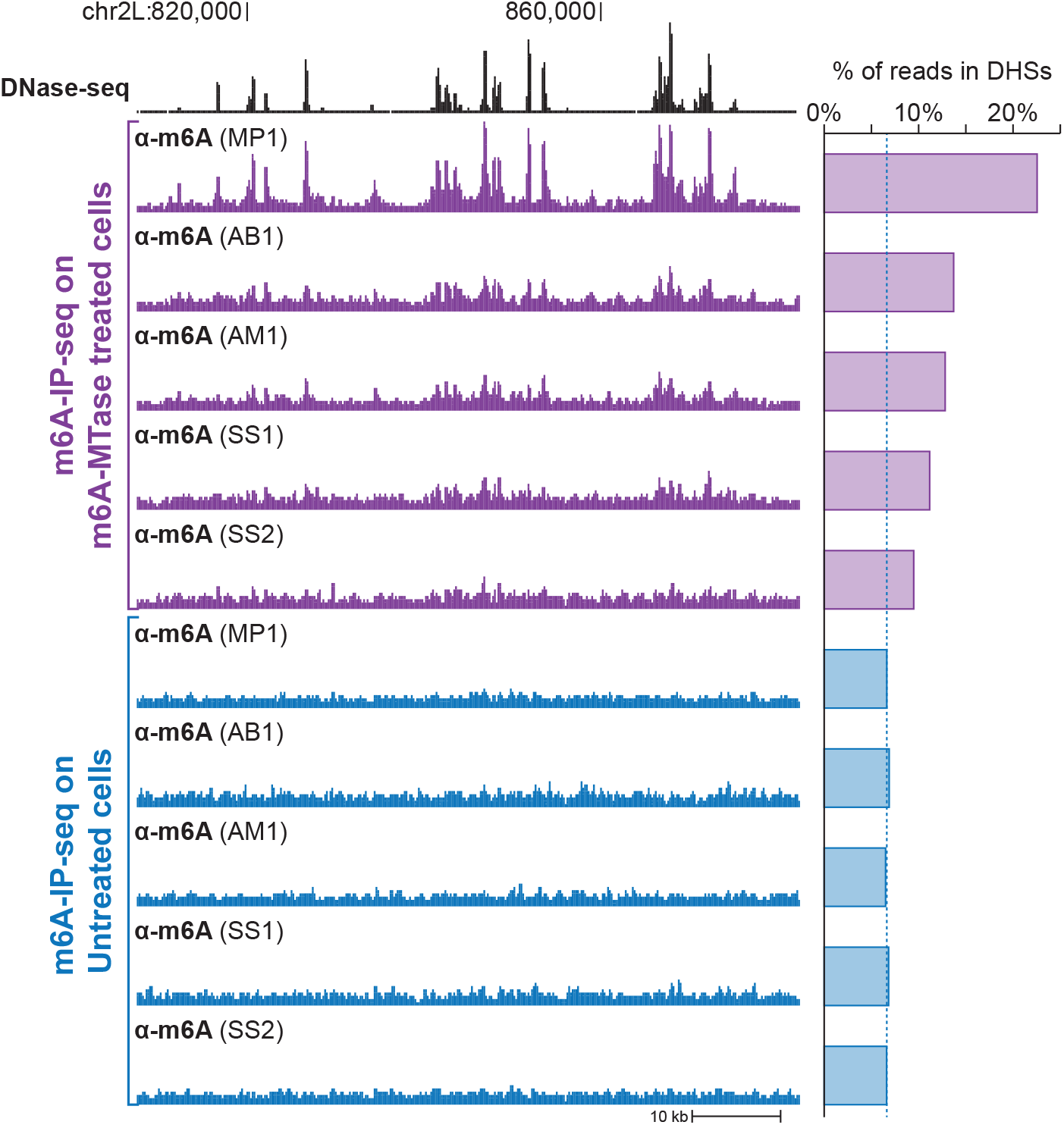
Detection of m6A-modified genomic loci using anti-DNA-m6A antibodies. (left) Genomic locus comparing the relationship between DNasel-seq and m6A-IP-seq signal. m6A-IP-seq per-formed using 5 separate antibodies on untreated S2 cell genomic DNA, or genomic DNA from S2 cell nu-clei treated with 0.6 U of a non-specific m6A-MTase. Y-axis is identical for all m6A-IP-seq experiments. (right) Percentage of sequencing reads that are localized to DNaseI hypersensitive sites for each of the m6A-IP-seq experiments. Higher values are indicative of higher antibody specificity.

**Supplementary Figure 4.**
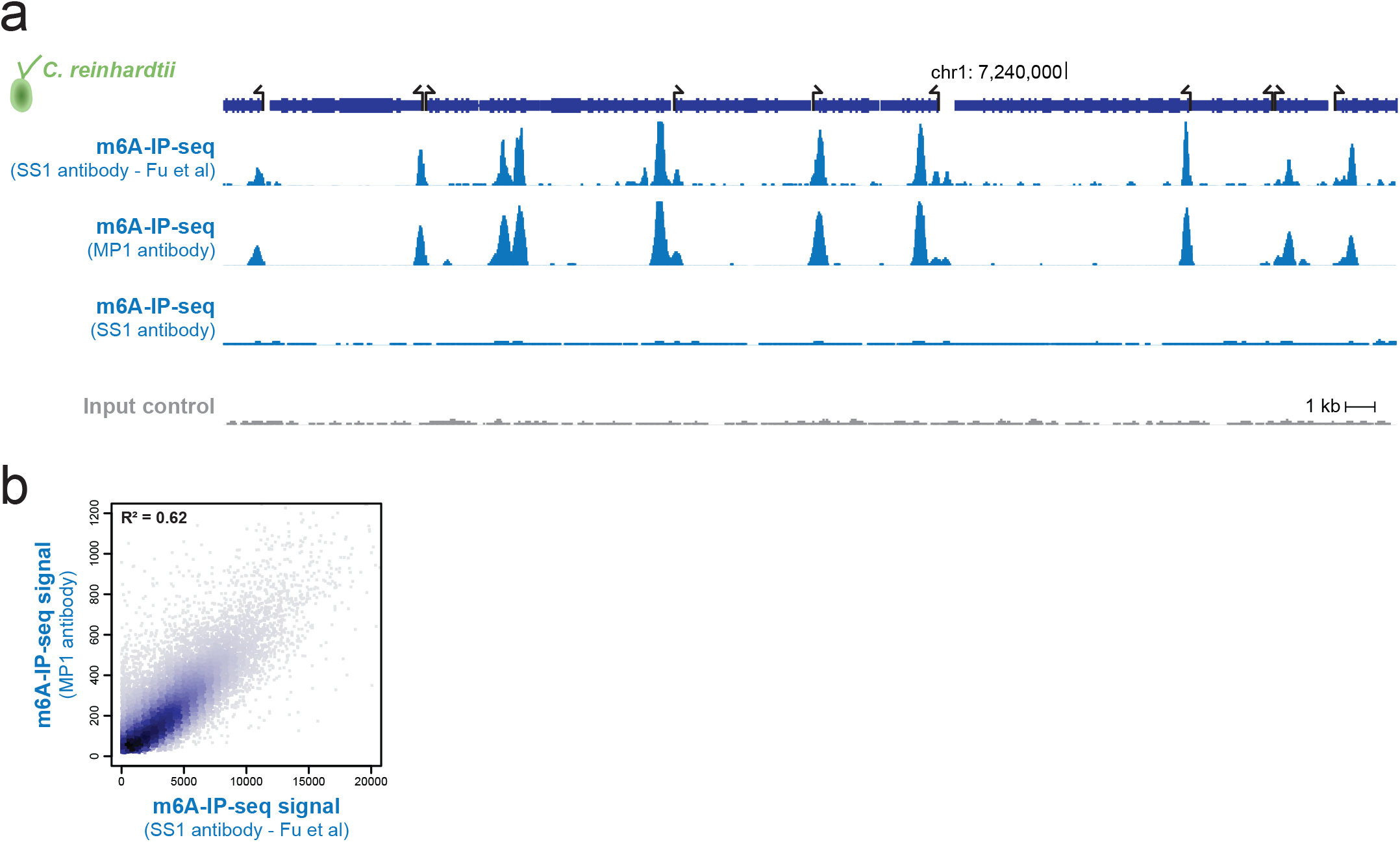
Site-specific DNA-m6A methylation in *C. reinhardtii*. (a) Genomic locus from *C. reinhardtii* comparing the relationship between m6A-IP-seq signal collected using either the MP1 or SS1 antibody, m6A-IP-seq signal previously published using the SS1 antibody, and input control signal. Y-axis is identical for all experiments except the previously published track. (b) Density scatter plot showing the relationship between m6A-IP-seq signal collected using the MP1 antibody, and m6A-IP-seq signal previously published using the SS1 antibody.

**Supplementary Figure 5.**
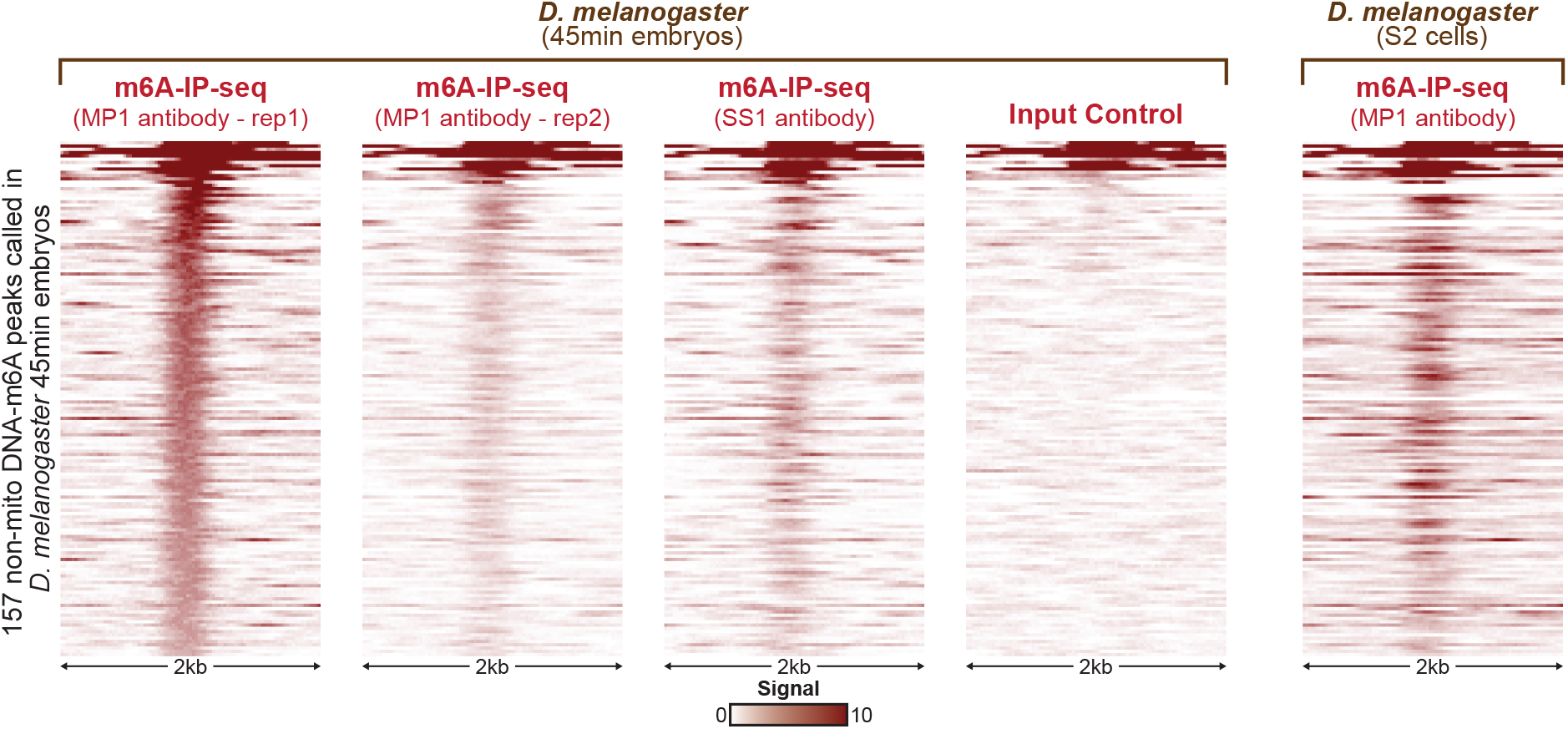
*D. melanogaster* 45-min embryos lack appreciable cell-selective site-specific DNA-m6A. Plots showing the signal strength surrounding each of the 157 non-mitochondrial DNA-m6A peaks identified in *D. melanogaster* 45-min embryos. Specifically, displayed is the m6A-IP-seq signal using two replicates of the MP1 antibody, as well as a single replicate of the SS1 antibody, and input control. Signal from m6A-IP-seq using the MP1 antibody on S2 cell genomic DNA is also displayed for comparison. The same scale is used for each of the five plots.

**Supplementary Figure 6.**
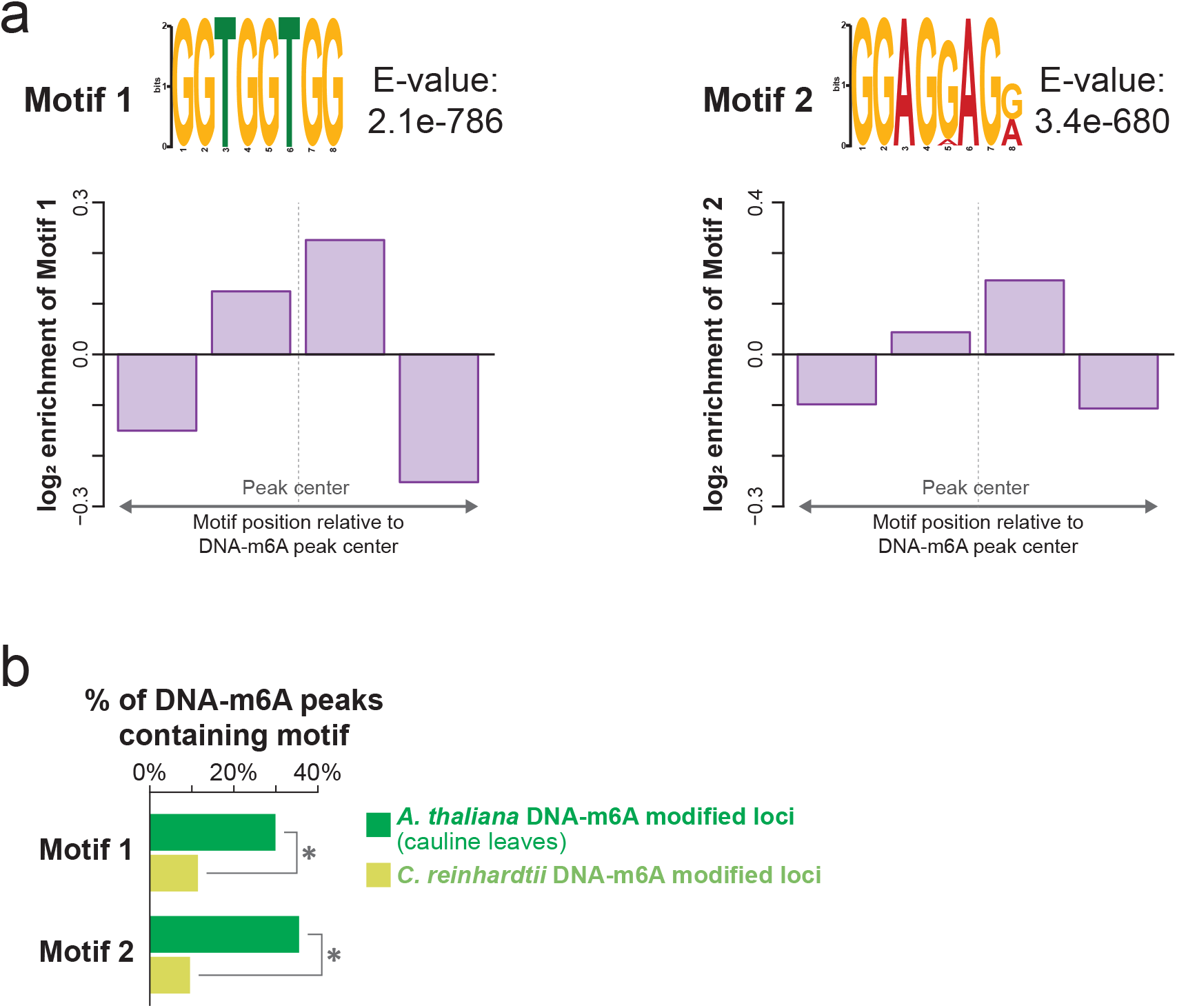
Enriched motifs in *A. thaliana* cauline leaf DNA-m6A peaks. (a) (top) Sequence logo and E-value for the top 2 motifs identified as being enriched in *A. thaliana* cauline leaf DNA-m6A peaks using MEME. (bottom) The distribution of each motif within *A. thaliana* cauline leaf DNA-m6A peaks relative to the size of each peak. Enrichment calculated based on the average density of each motif within *A. thaliana* cauline leaf DNA-m6A peaks. (b) The percentage of *A. thaliana* cauline leaf and *C. reinhardtii* DNA-m6A peaks that contain each of the motifs identified in panel a. * p-value <0.01 (z-test).

